# DynamicME: Dynamic simulation and refinement of integrated models of metabolism and protein expression

**DOI:** 10.1101/319962

**Authors:** Laurence Yang, Ali Ebrahim, Colton J. Lloyd, Michael A. Saunders, Bernhard O. Palsson

**Author notes:** Corresponding author at: Department of Bioengineering, University of California at San Diego, La Jolla, CA 92093, USA.

## Abstract

Genome-scale models of metabolism and macromolecular expression (ME models) enable systems-level computation of proteome allocation coupled to metabolic phenotype. We develop dynamicME, an algorithm enabling time-course simulation of cell metabolism and protein expression. Our dynamicME correctly predicted the substrate utilization hierarchy on mixed carbon substrate medium. We also found good agreement between predicted and measured time-course expression profiles. ME models involve considerably more parameters than metabolic models (M models). We thus present two methods to calibrate ME models, specifically using time-course measurements such as from a (fed-) batch culture. Overall, dynamicME and the methods presented provide novel methods for understanding proteome allocation and metabolism under complex and transient environments, and to utilize time-course cell culture data for model-based interpretation or model refinement.

## 1. Introduction

Computing the genotype-phenotype relationship is a fundamental challenge for computational biologists. Constraint-based reconstruction and analysis (COBRA) provides one approach for systems-level computation of biological networks using genome-scale biochemical network reconstructions [1]. Flux Balance Analysis (FBA) [2] in particular simulates flux distributions through a metabolic network by optimizing a cellular objective, such as maximizing growth rate subject to physicochemical, regulatory and environmental constraints. COBRA has been used to address a large variety of biological problems [3], and many algorithmic extensions have been developed [4].

In an important extension of FBA (FBAwMC), the hierarchy of substrate utilization in mixed carbon media was predicted correctly by imposing intracellular macromolecule crowding constraints [5]. The constraints imposed were based on approximate crowding coefficients for cytosolic enzymes based on estimated molar volume and catalytic efficiency.

Recently, genome-scale reconstructions have expanded significantly with development of integrated models of metabolism and macromolecular expression (ME models) [6, 7, 8, 9, 10]. ME models explicitly compute transcription and translation machinery requirements to support metabolic flux distributions. The latest *E. coli* ME models [9, 10] account for 80% of the proteome by mass and predict the allocation and limitation of proteome toward cellular functions during optimal growth [11]. Therefore, ME models considerably expand the scope of systems-level investigation and computation across multiple biological scales and processes.

In this work, we extend dynamic flux balance analysis (dFBA) [12, 13] to the time-course simulation of metabolism and proteome allocation using ME models [9, 10]. Recent algorithmic and modeling advances [14] for ME models have been critical for enabling this procedure, which we call dynamicME. We use dynamicME to explore the substrate utilization hierarchy of mixed substrate media from an optimal proteome allocation perspective. We also develop two approaches for refining ME model parameters using time-course metabolite data, which are often available from cell culture experiments. Thus, dynamicME provides both fundamental and practical advances for computing and understanding cellular metabolism and proteome allocation at the genome-scale.

## 2. Methods

### 2.1. Dynamic ME simulation

Our dynamicME implementation extends dynamic FBA (dFBA) [12, 13], which was developed for metabolic models (M-models). Because ME-models are considerably more computationally intensive than M-models, we made several simplifications and modifications to dFBA. First, the uptake rate of a substrate was not made a function of its extracellular concentration. Instead, flux bounds were set to zero if the substrate was depleted (zero concentration), or to a finite value otherwise.

This simplification allowed us to make the next simplification. That is, simulations were not re-performed at every time-step. Instead, once exchange (i.e., uptake and secretion) fluxes were computed by the ME-model, the same fluxes were used to compute the extracellular metabolite concentration profile over subsequent time-steps. At each time-step, dynamicME checked whether a substrate became depleted (fully consumed) or newly available, e.g., by feeding for a fed-batch process or secretion of re-consumable metabolites. If so, a new ME computation was performed with the updated exchange flux bounds. The exchange fluxes and growth rate were then updated according to the new optimal solution. These updated values were used to compute biomass and metabolite concentrations. This procedure was repeated until the batch time was reached. We note that concentration-dependent uptake rates or different feed schedules can still be implemented, albeit with added computational cost. The overall dynamicME procedure is shown in Fig. 1.

**Figure 1:**
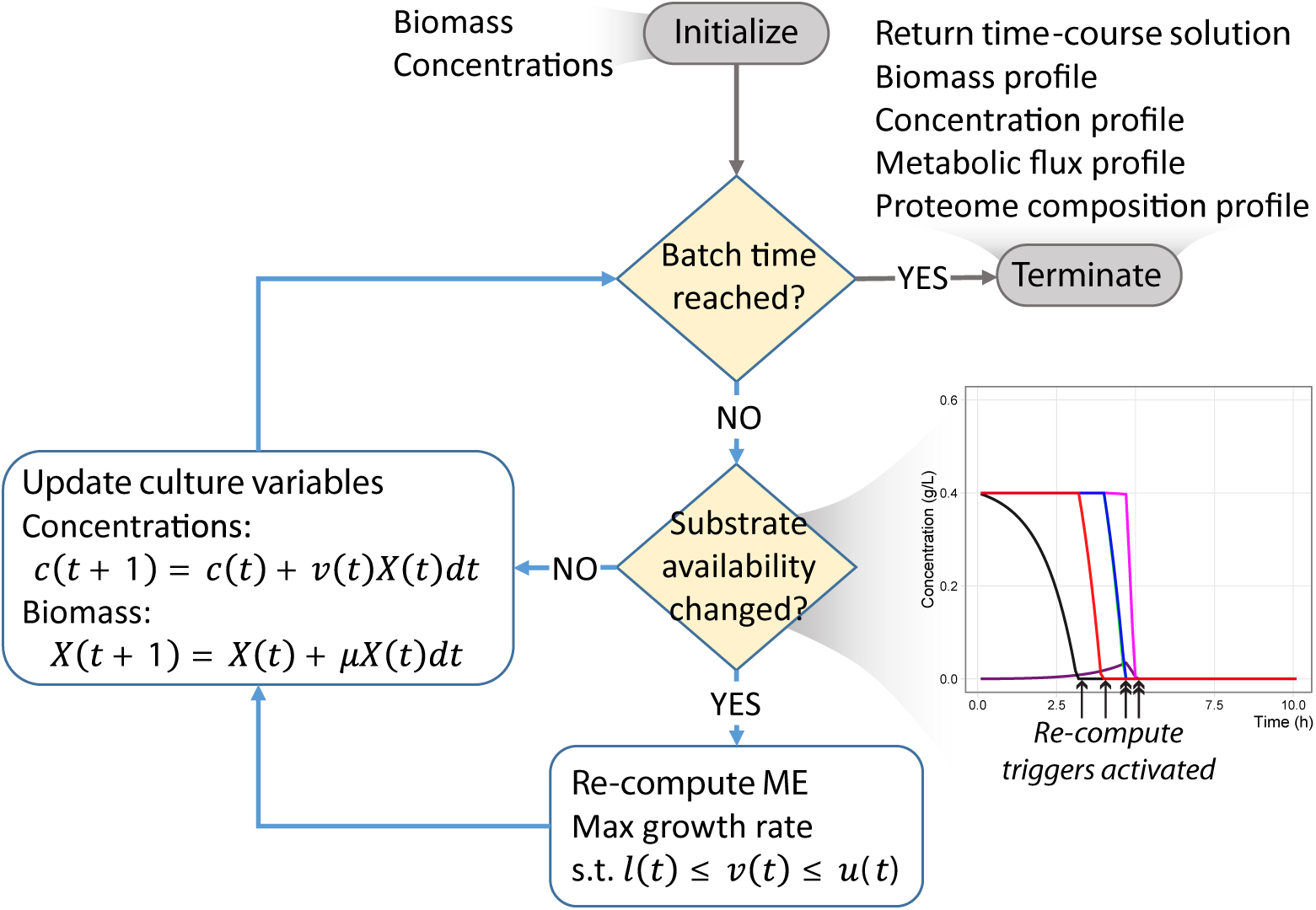
Schematic of the DynamicME procedure. The culture is divided into smaller time-steps and extracellular concentrations and biomass are updated at each timepoint. Metabolite exchange fluxes are computed whenever substrate availability changes due to metabolite depletion, feed, or secretion. With each ME simulation, the metabolic flux distribution and proteome composition are also updated.

### 2.2. Growth maximization for ME models

To maximize growth rate at each time-step, we used the solveME python module [14]. Specifically, we used bisection (binary search) as in [9] to maximize growth rate to six decimal points. SolveME uses the 128-bit (quad-precision) LP and NLP solver Quad MINOS 5.6 (qMINOS) [15, 16]. All qMINOS runs were performed with feasibility and optimality tolerances of 10^−20^. These tight tolerances were used to capture solutions involving fluxes as small as 10^−16^ mmol/gDW/h and were made possible through the quad-precision capabilities of qMINOS. The model used for this study are available on Github [17]. The COBRAme software [18] was used for building and developing the ME model.

### 2.3. Model calibration using literature data

A number of adaptive laboratory evolution (ALE) studies have now demonstrated that the proteome of wild-type *E. coli* is not optimally allocated or efficient for every single nutrient. To accurately reflect this wild-type proteome state, we calibrated the model with respect to several known enzymatic limitations of wild-type *E. coli.* First, glycerol kinase is known to be significantly less efficient for wild-type compared to ALE endpoints [19]. Second, ALE on lactate minimal medium showed multiple limitations in lactate utilization and enzymes near the phosphoenolpyruvate (PEP) node [20]. Third, respiration is known to have higher proteomic cost than fermentation, leading to acetate overflow [21]. Based on these observations, we calibrated the corresponding model parameters (i.e., effective rate constants—see section below for details). For example, we imposed a realistic turnover rate for isocitrate dehydrogenase based on literature data, effectively increasing proteomic cost for respiration. All calibrated parameters are listed in Table S1 along with their original and adjusted values.

### 2.4. Sensitivity analysis of dynamic simulations

In the ME model, effective rate constants (*k_eff_*) relate metabolic flux v to enzyme concentration by the relationship *v* = *k*_eff_ · *e*, where *e* is the enzyme concentration [9]. Precise estimates for these parameters are not available for many reactions and enzymes; therefore, an important step in ME model-based studies has been to assess sensitivity of predictions to these uncertainties [22]. In this study, we investigated the sensitivity of dynamicME predictions to uncertainties in *k*_eff_. We perturbed *k*_eff_ values from 0.1 to 10 times the nominal values. To avoid exploring the full parameter space consisting of thousands of keff values, we chose relevant pathways and perturbed only these reactions (Table S1). We generated 200 random samples. Perturbed ME models having good fit to measured metabolite concentration profiles were treated as an ensemble. The exact determination of ensembles is described below.

### 2.5. Archetypal analysis and ensemble of models

Archetypal analysis [23, 24] is a dimension-reduction method in which any data point is approximated as a convex combination of the computed archetypes; in turn, each archetype is a convex combination of the data points [23]. Each archetype lies on the convex hull of the data and represents a “pure” phenotype. In our study, we performed archetypal analysis on randomly perturbed samples of model-predicted time-course metabolite concentration profiles (Fig. 2). Thus, each archetype represents a distinct phenotype with a particular substrate utilization hierarchy.

**Figure 2:**
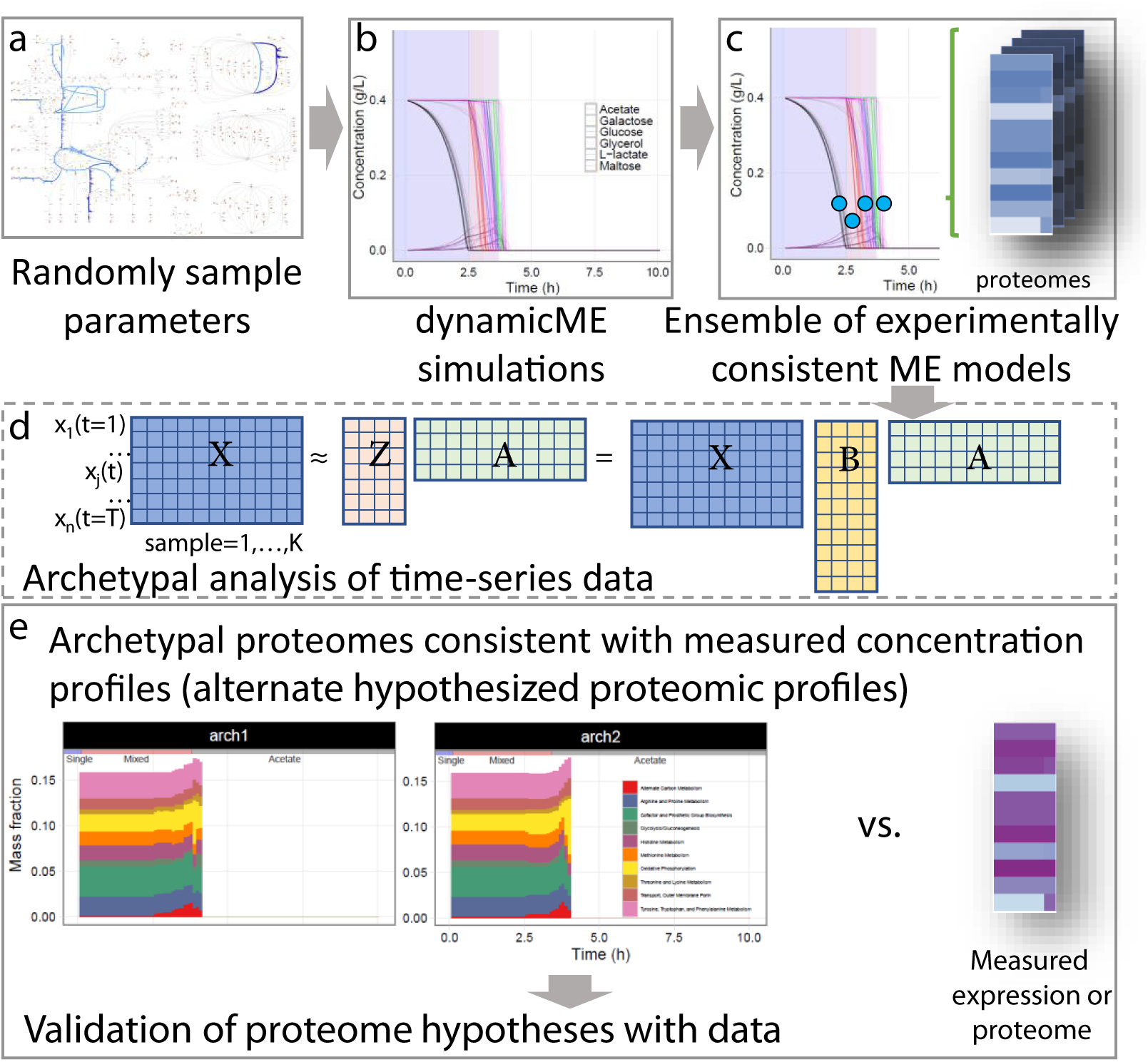
Ensemble model estimation and characterization procedure. (A-C) An ensemble of models with prediction error within a threshold was found by perturbing model parameters (*k*_eff_). (D) Archetypal analysis was used to characterize the ensemble where any data point is approximated as a convex combination of archetypes. (E) The predicted time-course proteome profile from the ensemble with best fit to measured metabolite concentrations was compared with measured time-course expression profiles.

To prepare data for archetypal analysis, timepoints and metabolites were collapsed, resulting in a 2D matrix of features (timepoints-and-metabolites) × samples. Given the archetypal matrix *Z*, original (collapsed) data matrix *X*, and the matrix *B*, *Z* = *XB* (Fig. 2d), where *B_ij_ ≥* 0 and 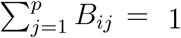 for *p* archetypes; therefore, the archetypes are constrained to be convex combinations of the data points *X*. Once archetypes were determined, the proteome and exchange flux time-series profiles were also mapped to the archetypes using *B*. The best number of archetypes was chosen using the elbow method from a scree plot [25] (Fig. S1). Archetypal analysis was performed using the `spams` Python module [26].

### 2.6. Optimal parameter estimation via meta-heuristic optimization

We developed an optimization-based procedure to match time-course concentration profiles by estimating *k*_eff_ values. For optimization, we used a gradient-free meta-heuristic method (list-based threshold accepting) [27] because of its efficiency and flexibility. We developed a parallel implementation of this optimization method for increased efficiency (Fig. S2). The implementation allows each parallel node (CPU thread) to choose between following its local search trajectory or restarting the search from the current best solution. This parallel communication was implemented using MPI via the mpi4py Python module [28]. The objective function was the sum of squared errors between measured and predicted extracellular metabolite concentration profiles.

### 2.7. Model validation using time-series expression profiles

To validate proteome allocation predictions, we computed the time-lagged cross-correlation between simulated and measured time-course proteome profiles. Lagged cross-correlation measures the similarity between two time-series where one lags the other, and has been particularly useful for analyzing time-course expression profiles. For example, it was used to study regulatory interactions of galactose metabolism in *E. coli* [29]. To compute lagged cross-correlation we used the `R` function `ccf` [30].

## 3. Results and Discussion

### 3.1. Growth on mixed substrates

When grown on complex media, *E. coli* uses substrates preferentially or simultaneously, depending on growth conditions [5]. Without additional constraints, FBA may erroneously predict simultaneous uptake of all substrates [5]. FBA with molecular crowding (FBAwMC) improves FBA by adding molecular crowding constraints, and correctly predicted substrate utilization hierarchy under a five-carbon medium [5].

We hypothesized that proteome-limited cellular growth would exhibit a hierarchy of preferential and simultaneous substrate utilization on mixed substrate media. To test this hypothesis, we implemented the dynamicME procedure: namely, time-course simulation of genome-scale integrated models of metabolism and macromolecular expression (ME-models) (Fig. 1). DynamicME extends dynamic FBA (dFBA) [13] to ME-models (see Methods).

Using dynamicME, we simulated cellular growth on the five-carbon mixed substrate media studied by Beg et al. [5] and the simulated metabolite concentration profiles were compared with measurements. To simulate growth on nutrient-excess batch culture, carbon substrate uptake rates were effectively unconstrained (i.e., lower bound = −1000 mmol/gDW/h). Therefore, total proteome limitation became the active constraint rather than nutrient limitation. Oxygen uptake rate was constrained to −20 mmol/gDW/h to reflect transport limitations that are independent of proteome limitation.

DynamicME correctly predicted the majority of substrate uptake hierarchy characteristics, including the single substrate utilization (glucose), mixed utilization, and acetate reconsumption phases observed by Beg et al. [5] (Fig. 3). We found a few differences between simulated and measured profiles. Overall, metabolites were consumed more rapidly than experimentally observed. Also, acetate secretion was lower than measured, and maltose was predicted to be utilized earlier than in experiments.

**Figure 3:**
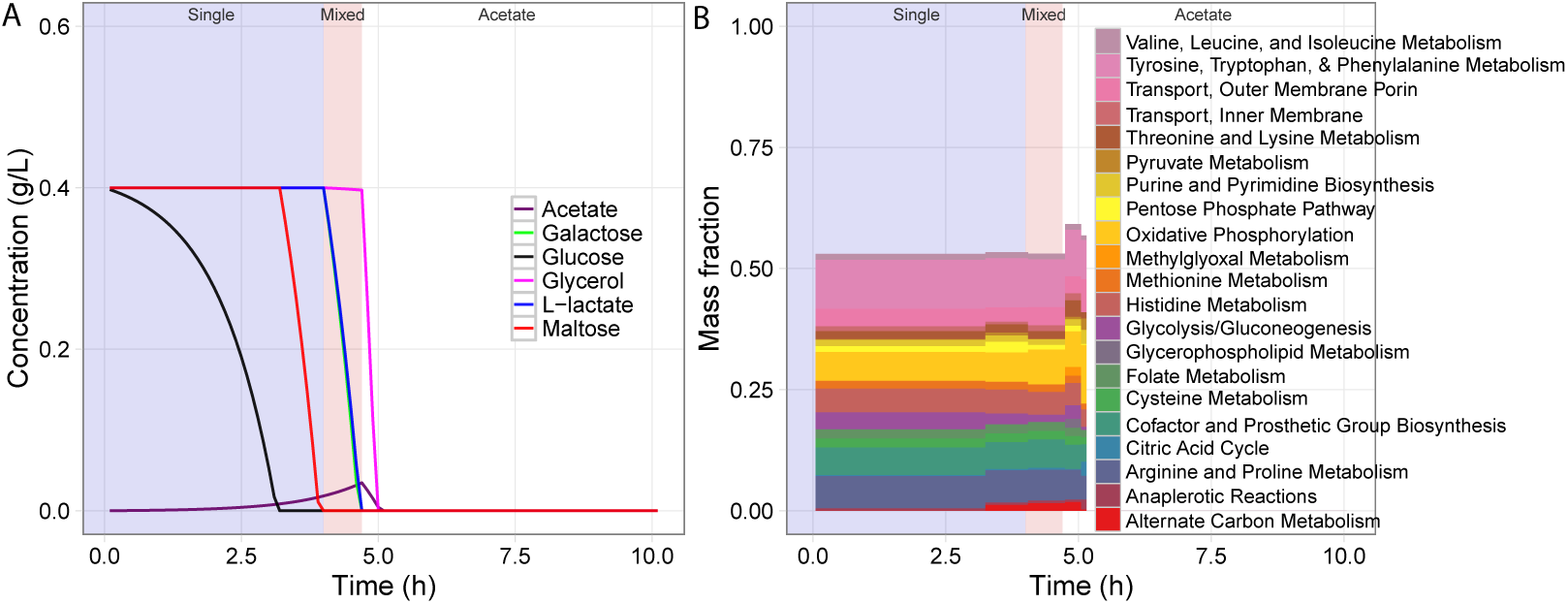
Predicted substrate uptake hierarchy and proteome allocation. (A) Predicted time-course metabolite concentration profile. (B) Predicted time-course proteome mass fraction allocation profile.

In the absence of additional constraints, FBA was shown to predict optimal states that accurately reflect ALE endpoints but may exceed the efficiency of wild-type cells [31, 32, 33, 34]. To account for this discrepancy, we implemented a model-calibration procedure to better reflect observed metabolic and expression profiles, as described in the following section.

### 3.2. Model calibration for experimentally consistent concentration time-course profiles

Both the rate and hierarchy of substrate utilization are affected by ME-model parameters. In particular, the effective rate constants *k*_eff_ influence predicted pathway usage [35, 36]. We thus investigated the sensitivity of predicted substrate utilization hierarchy to uncertainty in *k*_eff_ values.

First, we performed 200 random perturbations of *k*_eff_ values and performed dynamicME simulations for the perturbed models. The predictions showed large variations with respect to substrate utilization hierarchy. To aid interpretation, we performed archetypal analysis [23, 24] on the time-course metabolite concentration profiles and identified five archetypes (Fig. 4) as described in Methods. The five archetypes showed considerable variation in substrate utilization hierarchy, reflecting the sensitivity of predictions to uncertainty in *k*_eff_ values. Of the five archetypes, archetype 4 most closely resembled experiments (Fig. 4). The archetypal model correctly predicted the sequence of glucose uptake followed by mixed utilization of maltose, lactate, and galactose, and finally glycerol uptake and acetate re-consumption. The acetate secretion rate was also significantly higher than the initial model and matched measurements better.

**Figure 4:**
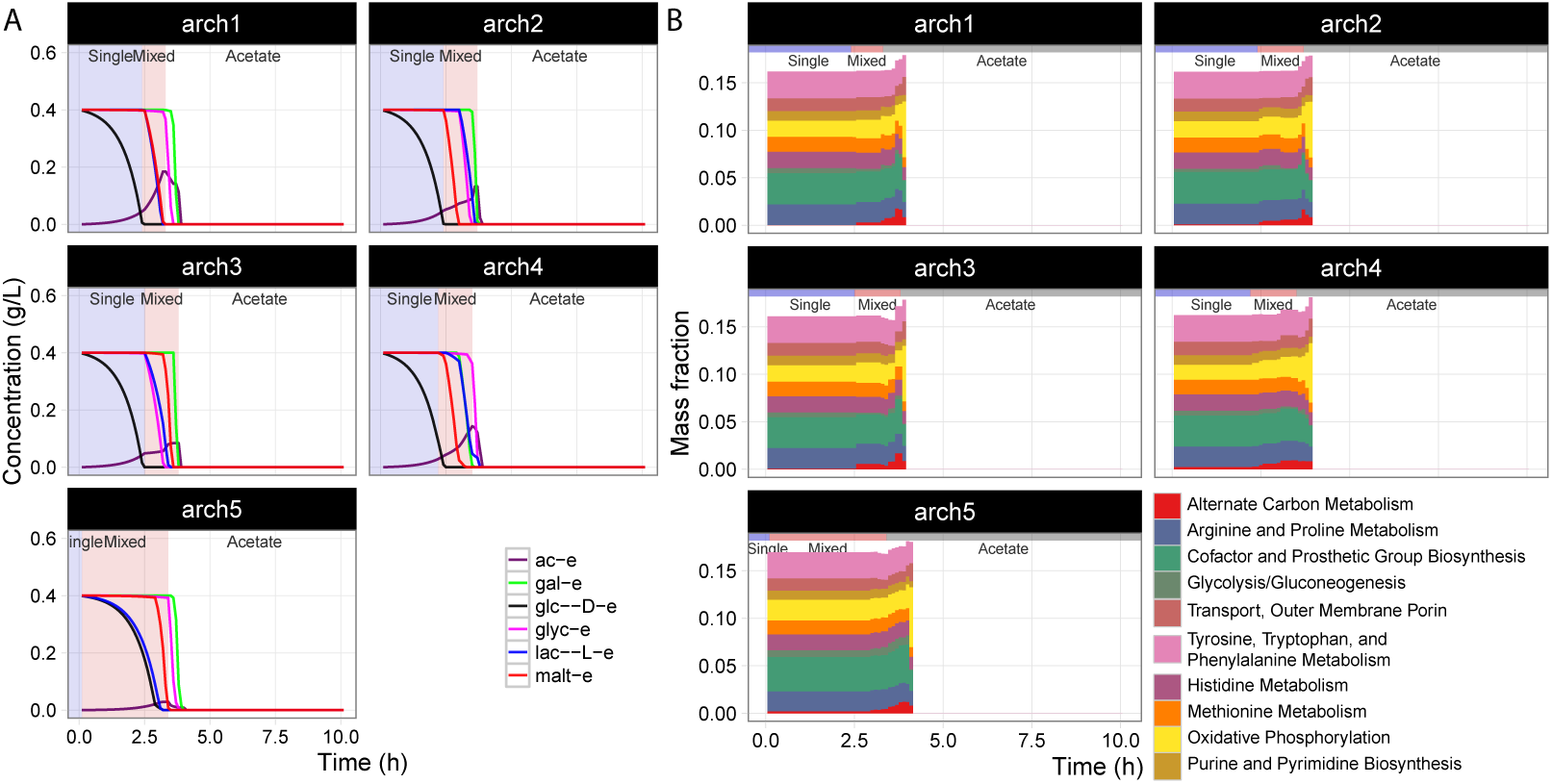
Sensitivity and archetypal analysis. Archetypal time-course concentration (A) and proteome mass fraction allocation (B) profiles were computed from simulations with 200 randomly perturbed *k*_eff_ parameters.

We also implemented an alternative approach to fit measured concentrations using meta-heuristic optimization (Fig. S2, Fig. S3). The optimal profiles were similar to that of archetype 4. Thus, we proceeded with subsequent analyses using archetype 4, which in turn represents an ensemble of experimentally-consistent ME models with differing parameter values.

### 3.3. Predicting time-course proteome allocation

An important novelty of dynamicME is explicit computation of proteome allocation over a time-course simulation. For the mixed substrate medium, dynamicME computed distinct proteome compositions over time, corresponding to the changing metabolic modes (Fig 3). We compared computed proteome dynamics with measured time-series microarray data [5]. For validation, we used the proteome profile from the most accurate archetype (archetype 4) as determined in the previous section (Fig. 4).

To validate proteome allocation predictions, we computed the lagged cross-correlation [37] between simulated and measured time-course proteome profiles. The median lagged cross-correlation was 0.67 with values between −0.50 to 0.92 (Fig 5A). Consistent with metabolite concentration profiles, simulated proteome dynamics were faster than measured, with a median lag of 0.8 hours (Fig 5A).

**Figure 5:**
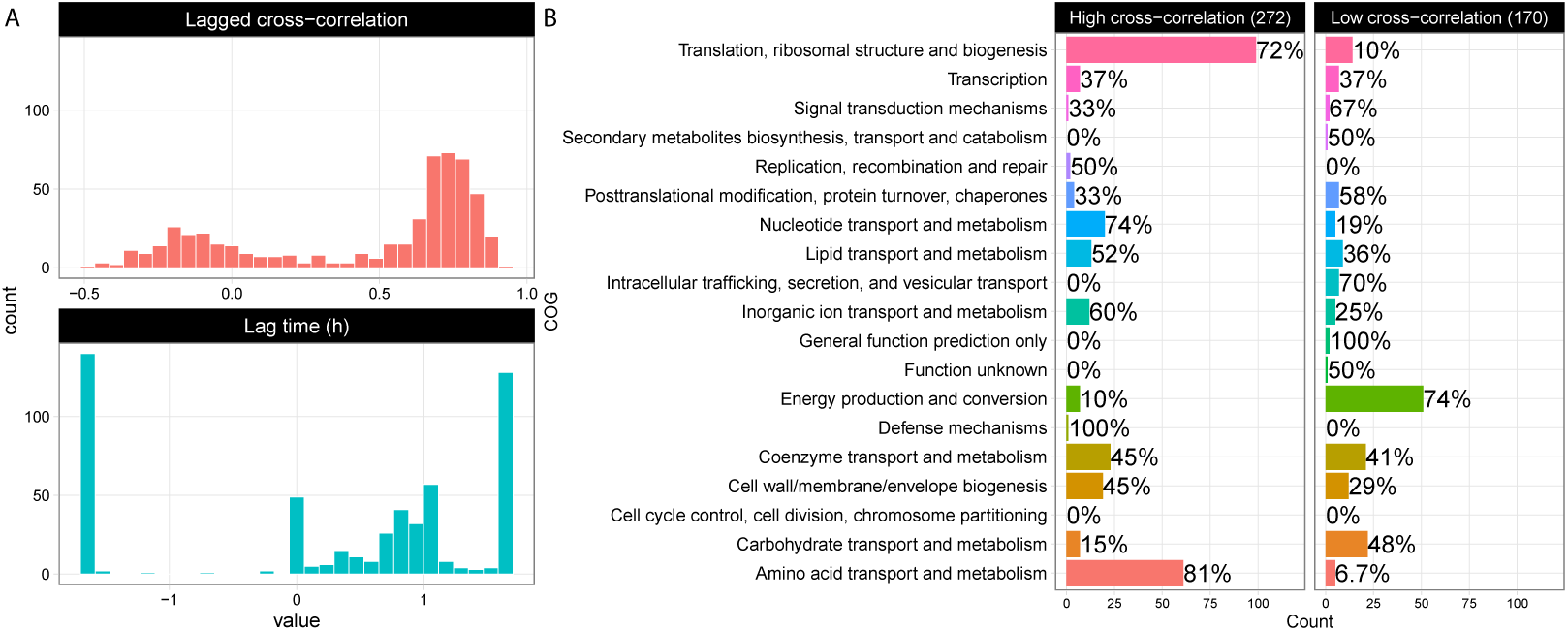
Lagged cross-correlation of simulated vs. measured expression. (A) Histograms of lagged cross-correlation values and lag time. (B) Functional groups (COGs) of genes with lagged cross-correlations that were low (below 0.25) or high (above 0.67, the median).

In addition, certain functional gene sets were predicted better than others. For example, of 138 genes in the COG “Translation, ribosomal structure and biogenesis” dynamicME predicted 72% with high (above 0.67, the median) and 10% with low (below 0.25) cross-correlation, respectively (Fig 5B). Similarly, “nucleotide transport and metabolism,” “amino acid transport and metabolism,” and “inorganic ion transport and metabolism” were predicted with high cross-correlation. In contrast, of 69 “Energy production and conversion” genes, 74% had low cross-correlation.

## 4. Conclusions

In this study we developed dynamicME, an algorithm for simulating time-course metabolic and proteomic profiles using genome-scale models of metabolism and macromolecular expression (ME-models). We found that dynamicME correctly predicted substrate utilization hierarchy under a five-carbon mixed substrate medium. The biological basis for this hierarchy was proteome-limited cellular growth.

To account for the tendency of constraint-based models, including ME-models, to over-predict metabolic efficiency over wild-type cells, as well as parameter uncertainty, we implemented a model calibration procedure. In this study we focused on perturbing the effective rate constants (*k*_eff_) to match metabolite concentration profiles better. We arrived at a set of models showing improved prediction of the substrate utilization hierarchy. We note that sensitivity of ME model predictions to *k*_eff_ values has been investigated in several studies including a non-dynamic context [35], in relation to expression of protein groups (or sectors) [38], and for defining a core proteome [36]. However, the sensitivity of the predicted sequence and preference of utilizing mixed carbon substrates over time had not been investigated prior to the present study.

A notable feature of DynamicME is its ability to predict time-course proteome allocation profiles. We observed good agreement between measured and computed time-course expression profiles (median lagged crosscorrelation of 0.67). Meanwhile, one subtle difference between measured and predicted time-course profiles was that measured profiles changed less abruptly due to process time constants of transcription and translation dynamics. We note that the original dFBA for metabolic models implemented rate-of-change constraints on fluxes precisely to account for this phenomenon. In future work, dynamicME could include a similar constraint, directly on the transcription and translation fluxes.

ME-models compute cellular resource allocation tradeoffs at the proteome scale [39]. This expanded biological scope and predictive capability of ME models is expected to become increasingly useful for biotechnological applications [40]. For example, the metabolic and proteomic burden to the host of expressing biochemical production pathways can be computed explicitly using ME-models. We have shown here that it is furthermore possible to compute how these genome-wide cellular resource dynamics determine transient shifts in metabolic modes under biotechnologically relevant culture conditions: complex media with transient substrate availability. Thus, as ME-models continue to be reconstructed for organisms of biotechnological importance, dynamicME will be a useful approach for analyzing physiological and omics data from cell culture, and for model-aided biotechnological applications that require robust cell factory operation under environmental fluctuations [41].

## Acknowledgments

This work was funded by the National Institute of General Medical Sciences of the National Institutes of Health [awards U01GM102098 and R01GM057089] and the Novo Nordisk Foundation through the Center for Biosustainability at the Technical University of Denmark (NNF10CC1016517). This research used resources of the National Energy Research Scientific Computing Center, which is supported by the Office of Science of the US Department of Energy under Contract No. DE-AC02-05CH11231. The content is solely the responsibility of the authors and does not necessarily represent the official views of the funding agencies.

## Supplementary material

**Fig. S1.**
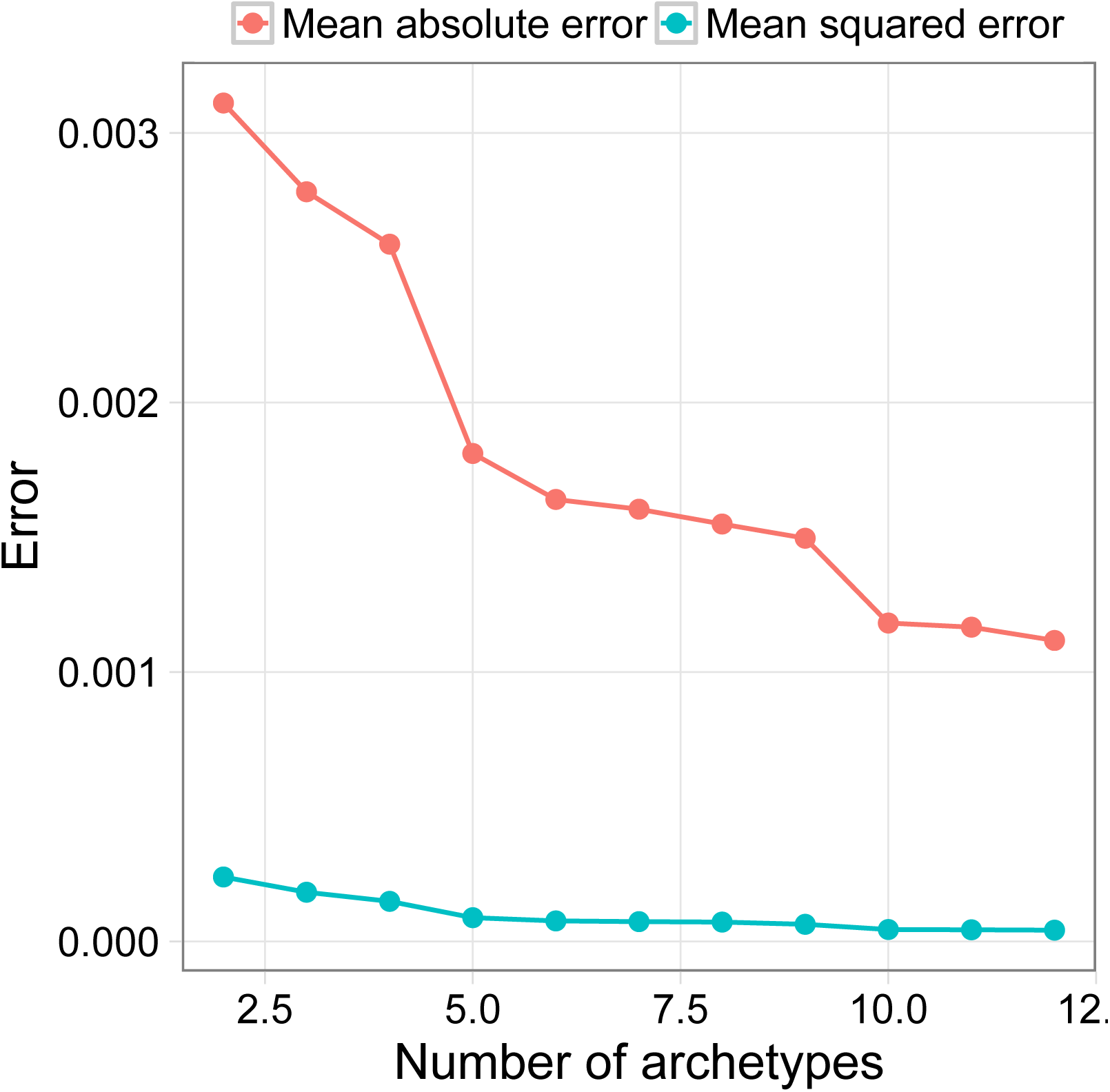
Scree plot for determining number of archetypes. A notable elbow is observed for five archetypes.

**Fig. S2.**
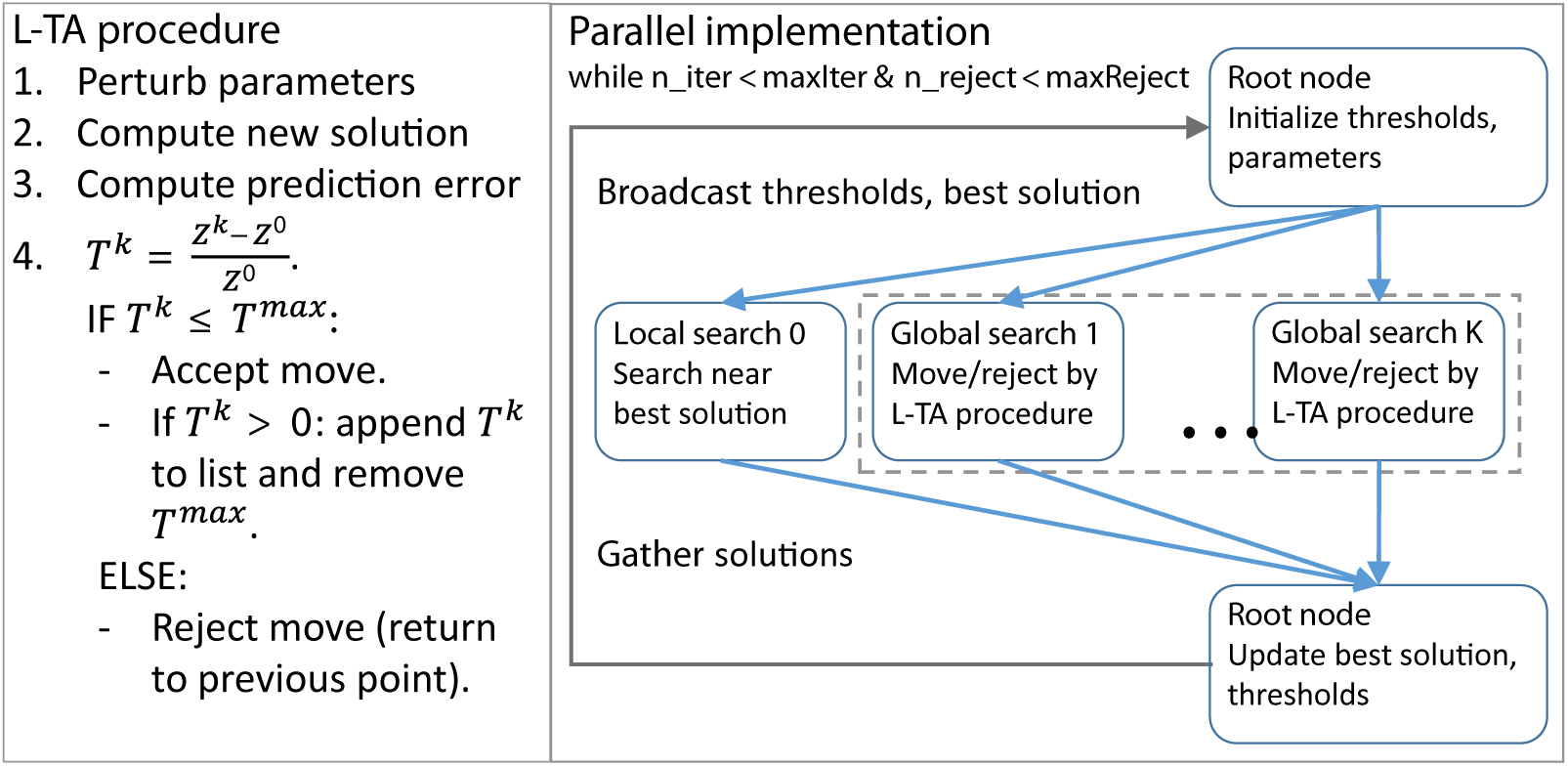
Parameter estimation procedure. We developed a parallel implementation of a meta-heuristic optimization procedure. L-TA: list-based threshold accepting algorithm [27].

**Fig. S3.**
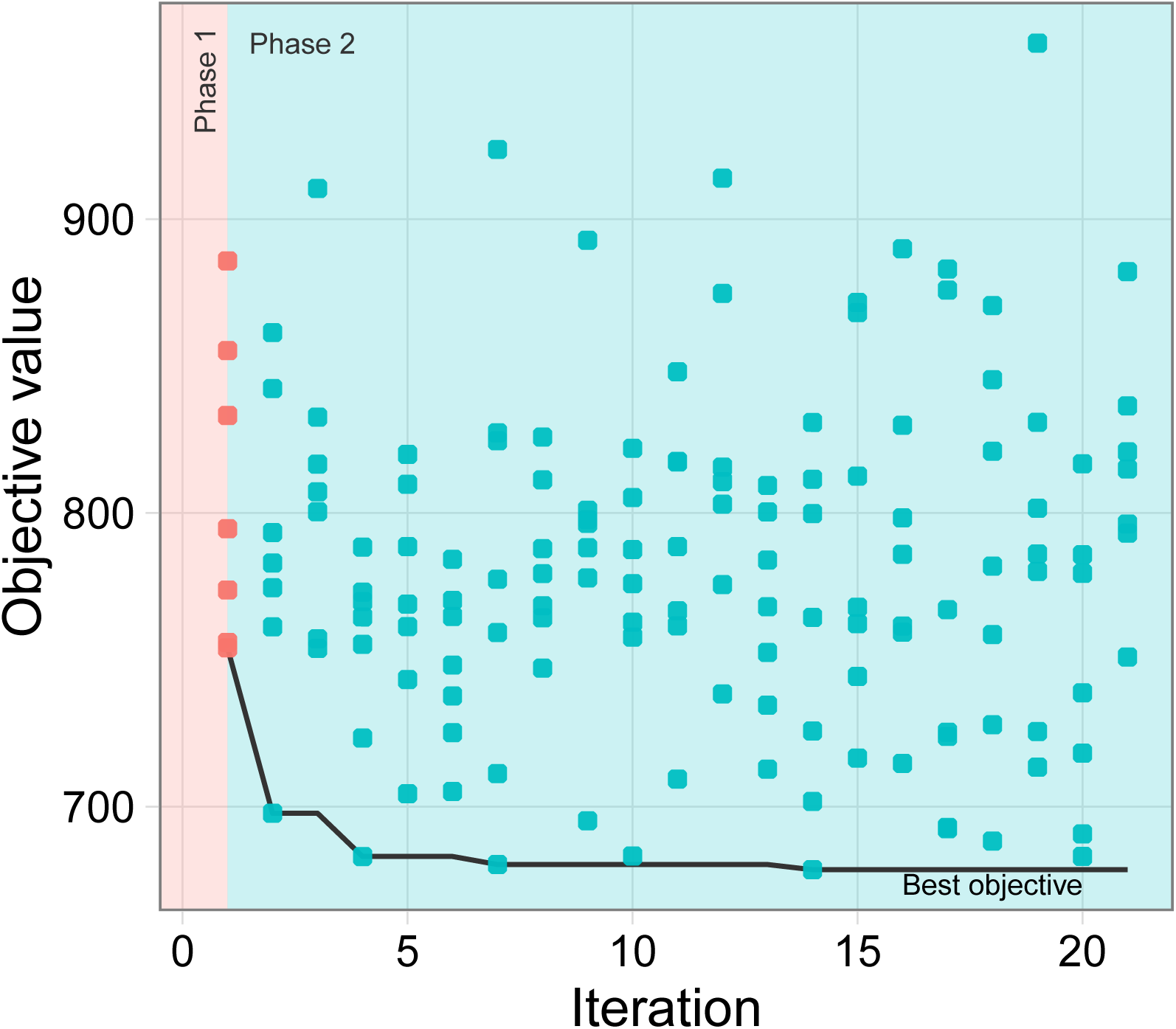
Parameter estimation results. The parallel L-TA optimization procedure successfully estimated model parameters that improved consistency with measured concentration profiles. Seven parallel nodes were used here: 1 local and 6 global nodes (see Fig. S2) for explanation of nodes.

**Table S1.** Reactions with perturbed. *k*_eff_. A subset of the genome-scale metabolic network was perturbed with respect to *k*_eff_ values, either manually or randomly.

